# Early Emergence of Auditory Quantity Discrimination in Domestic Chicks

**DOI:** 10.64898/2026.04.08.717196

**Authors:** Elena Eccher, Orsola Rosa Salva, Cinzia Chiandetti, Giorgio Vallortigara

## Abstract

Numerical abilities are widespread in the animal kingdom and are not exclusive to humans. Domestic chicks (Gallus gallus) have been shown to discriminate numerosities spontaneously, but prior research has focused exclusively on the visual modality. Whether chicks can discriminate numerical information in the auditory domain remains unknown, despite evidence that they can perceive other auditory features such as tone and rhythm. In this study, we investigated spontaneous numerical discrimination in the auditory modality in naïve domestic chicks. In Experiment 1, newly-hatched chicks were tested for their ability to discriminate between two auditory sequences differing in numerosity (4 vs. 12 identical sounds), with and without controlling for continuous variables such as duration and total sound amount. Experiment 2 examined chicks’ filial imprinting responses to familiar or unfamiliar numerosities. Experiment 3 controlled for potential spontaneous preferences for a single longer sound versus a shorter one. Our results showed a preference for the 12-sound sequence only when duration and total sound amount were not matched. When these continuous variables were controlled, no spontaneous numerical preference emerged. Experiment 2 revealed an overall preference for the 12-sound sequence regardless of imprinting conditions, while Experiment 3 confirmed that chicks do not have an inherent preference for longer sounds. These findings suggest that chicks are sensitive to overall magnitude in the auditory domain but do not spontaneously discriminate numerical differences when other continuous variables are held constant. Future studies will explore how specific stimulus features, such as heterogeneity of sounds, influence these preferences.

## INTRODUCTION

Early theories on the origins of numerical competence viewed words and symbols as essential prerequisites for understanding quantity and number, thereby attributing this capacity primarily to language (Bloom, 1994; Chomsky, 2006). More recent research, however, has demonstrated that the ability to discriminate and estimate quantities can emerge independently of linguistic or symbolic representations. Far from being unique to humans, such *Number Sense*, has been observed across a wide range of animal species, suggesting that it emerged early in evolution and serves important adaptive functions (Nieder, 2020): optimizing foraging and hunting strategies (Hanus & Call, 2007; Panteleeva et al., 2013), selecting mates (Kitchen, 2004; Lyon, 2003), and managing social interactions (Benson-Amram et al., 2018; Grinnell et al., 1995; Wilson et al., 2001). These numerical abilities rely on the Approximate Number System (ANS), a non-verbal, innate mechanism, shared across species, that supports rapid perception, representation, and manipulation of quantities without relying on symbolic representation (Feigenson et al., 2004). For instance, primates, such as rhesus macaques and chimpanzees, can match sets based on quantity, perform ordinal comparisons, and solve simple addition or subtraction tasks, both with visual arrays and sequential presentations (Anderson et al., 2007; Beran et al., 2019; Brannon & Terrace, 1998). These competences are not restricted to our closest relatives: other mammals, including elephants, dogs, and dolphins, discriminate quantities and make number-based decisions (Ward & Smuts, 2007; Perdue et al., 2012). Birds show particularly striking abilities, with corvids and domestic chicks both capable of discriminating small and large numerosities, generalising across contexts, and solving addition or subtraction problems (see Regolin et al., 2025 for a review). Even invertebrates, such as honeybees, have been shown to learn to associate numerosities with specific outcomes (Bortot et al., 2019). The ANS also shows similar properties in humans and other animals, operating in a ratio-dependent fashion consistent with Weber’s law, whereby discriminability depends on the proportional rather than the absolute difference between quantities (Gallistel & Gelman, 1992; Brannon, 2006; Nieder, 2019).

While research on humans helps to shed light on the most complex phenomena regarding numerical cognition (e.g., the neural basis supporting algebraic operations, Monti et al., 2012), comparative studies of animal cognition help to tackle challenging questions about the influence of innate mechanisms, culture, and language in the evolution of those same abilities (Platt & Spelke, 2009). Studying species that are evolutionarily distant from humans helps trace the phylogenetic history of the neural mechanisms underlying human cognition, shedding light on their ecological relevance and evolutionary origins. Moreover, investigating cognitive abilities in non-human animals allows us to observe how these abilities manifest in the absence of cultural influences. Because humans are highly immature at birth from a sensorimotor perspective, and their early sensory and social experiences are virtually impossible to control, it is extremely challenging to determine which cognitive abilities are truly present at birth. The same limitations apply to other altricial species, such as non-human primates (Sugita, 2008). In contrast, precocial species are born with advanced motor and sensory abilities, allowing them to stand, walk, and interact with their environment shortly after birth. Among them, domestic chicks (*Gallus gallus domesticus*) have become a key model for investigating innate predispositions and the foundations of numerical cognition, as they are fully developed at hatching and can be tested soon after birth, with sensory experiences precisely controlled thanks to their in ovo development (see Versace & Vallortigara, 2015).

A fundamental component of numerical competence is the ability to discriminate between different numerosities, often assessed through “larger versus smaller” comparisons (Vallortigara et al., 2010). Chicks reliably discriminate both small numerosities, such as 2 versus 3 items (Rugani et al., 2008; Rugani, Cavazzana, et al., 2013), and larger ones, such as 8 versus 12 items (Rugani et al., 2014; Rugani, Vallortigara, et al., 2013), in a ratio-dependent manner consistent with Weber’s law. They are also sensitive to proportional relations: it has been demonstrated that chicks can generalise across equal ratios, suggesting an early-emerging ability to encode and compare relational quantities (Rugani, Vallortigara, & Regolin, 2015; Rugani et al., 2016). Remarkably, very young chicks even showed arithmetic-like competencies. In a classic study, Rugani and colleagues (2009) reared chicks with five objects and, at test, presented them with occlusion scenarios where objects were hidden behind screens. Chicks consistently approached the panel, concealing the larger number of items, and, when objects were visibly transferred between the screens, they appeared to keep track of the addition or subtraction of items (Rugani et al., 2009). Such findings suggest that chicks can mentally represent changes in quantity over time (Rugani et al., 2011). In addition, they can encode ordinal information, identifying targets based on their position within a sequence (Rugani et al., 2007; Rugani & Regolin, 2020), even revealing spontaneous rightward biases in mapping numerosity into space (Rugani et al., 2010, 2020; Rugani, Vallortigara, Priftis, et al., 2015; Rugani & Regolin, 2021), akin to the Mental Number Line in humans (see Eccher et al., 2025 for a review). Recent evidence also shows that the chick brain contains neurons selectively tuned to specific numerosities (“number neurons”; Kobylkov et al., 2022), providing a potential neural basis for numerical representation similar to that observed in mammals (Nieder, 2016).

Often, chicks’ numerical competencies emerge without explicit training and can be revealed by their spontaneous choices or through filial imprinting. This is a form of learning by exposure, in which newly hatched chicks rapidly form attachments to visual or social stimuli (Bolhuis, 1991). Filial imprinting has been exploited to examine whether chicks can form attachments to a specific numerosity by rearing them with sets containing different numbers of visible objects (e.g., 1 vs. 3) and subsequently testing their preference for various numerosities. This has revealed that chicks can show different preferences, either for larger or familiar numerosity, depending on specific task features (Rugani et al., 2010)

Although a substantial body of research has examined numerical abilities in domestic chicks, these studies have exclusively relied on visual stimuli. Evidence from other species, however, indicates that numerical processing is not limited to vision but also extends to audition. For example, rats trained with three bursts of white noise responded most strongly to that numerosity, showing reduced responses to two or four bursts—suggesting genuine number discrimination rather than a simple “many-versus-few” distinction (Davis & Albert, 1986). In ecologically relevant situations, acoustic quantity estimation can also guide behaviour: lions, for instance, adjust their aggressive strategies depending on the number of roars heard from a rival group, effectively using acoustic input as a proxy for numerical estimation (McComb et al., 1994). Similarly, untrained cotton-top tamarin monkeys (*Saguinus oedipus*) can discriminate sequences of syllables based on the number of items, even when non-numerical cues such as duration, inter-stimulus interval, and overall energy were controlled for (Hauser et al., 2003). Interestingly, their performance revealed a clear ratio effect, strikingly similar to that observed in 9-month-old human infants when discriminating auditory sequences (Lipton & Spelke, 2004).

Chicks are capable of complex auditory processing: they can be imprinted prenatally on specific tones, and later recognise them after hatching (Grier et al., 1967), they show spontaneous preferences for consonant over dissonant melodies (Chiandetti & Vallortigara, 2011), and they are sensitive to temporal structure in sound, including rhythmic regularities and changes in auditory patterns (De Tommaso et al., 2019). Despite this, no study has directly tested auditory numerical discrimination in chicks. Addressing this question is essential in determining whether numerical cognition in this precocial species is truly modality-independent. In this study, we aimed to investigate newborn chicks’ ability to discriminate auditory numerosities using two methodological approaches commonly employed in chick research: spontaneous choice paradigms and filial imprinting. Here, we combine these two approaches to specifically test whether newborn chicks can spontaneously discriminate auditory sequences of different numerosities and whether they can acquire numerical information through imprinting.

## GENERAL METHODS

### Ethical Note

In accordance with Italian and European legislation, all experimental procedures included in this work were reviewed and approved by the Committee for Animal Welfare of the University of Trento. The experimental protocols were subsequently evaluated by the Italian National Institute of Health and authorised by the Italian Ministry of Health, Department of Food, Nutrition, and Public and Veterinary Health (permit number 987/2017). All procedures were carried out in full compliance with the relevant national and European regulations and guidelines.

### Animals

Fertilised eggs were supplied by a commercial hatchery (CRESCENTI Società Agricola S.r.l.—Allevamento Trepola—cod. Allevamento127BS105/2, Italy). Upon arrival, eggs were placed in an incubator (FIEM, MG 200/300 super rural) and maintained under standard incubation conditions (temperature: 37.7 °C; relative humidity: 40%) until embryonic day 19. At this point, eggs were transferred to a hatching unit (FIEM, MG 140/200 rurale) with the same temperature but increased humidity (60%). Both the incubators and the hatching room were kept in complete darkness. After data collection, chicks were given to local farmers. Sex determination was performed immediately following testing, using rapid inspection of wing feathers (a method applicable to this auto-sexing breed), carried out by experienced experimenters. The required sample size for each experiment was estimated through an a priori power analysis conducted with G*Power 3.1 (Faul et al., 2009), assuming an effect size of Cohen’s d = 0.35, a significance level of α = 0.05, and a desired power of 1 − β = 0.80. The analysis indicated that a minimum of 67 animals per experimental group would be required for a two-tailed one-sample Student’s t-test.This number was rounded up to 68 per condition to allow balanced left/right stimulus presentation across individuals. When two conditions were tested (Experiments 1a, 1b, and 2), this resulted in a total sample of 136 animals per experiment. Experiment 3 included only one condition and therefore had a sample of 68 animals.

### Apparatus

A wooden Y-shaped apparatus was used in all experiments, with the inner walls covered in sound-absorbing material to minimise reflections. At the end of each arm, a speaker (Z130

Stereo Speakers) was mounted at the chick’s height, approximately 10 cm from the floor. A task-irrelevant red object was hung in front of each choice area to encourage approach. The presence of an identical, salient visual cue on both sides helps motivate chicks to enter the choice zones while avoiding any directional bias. The objects were cylindrical (2 cm diameter

× 7 cm height) and were suspended 10 cm above the floor of the apparatus. Two LED strips were mounted above the two ends of the apparatus to illuminate its interior. No other light sources were present in the testing room. On the floor, a fine pencil line delimited the choice areas in the two arms of the Y-maze (20 cm × 35 cm), while the central section (20 cm × 20 cm) functioned as the no-choice area and starting point for the chicks. A webcam (Logitech, C922 Pro, HD stream webcam) positioned above the apparatus recorded an aerial view of the chicks’ movements throughout the test.

In all experiments, acoustic stimuli consisted of multiple repetitions of identical notes/tones (as described in each experiment section). Each sequence has been generated in version

2.0.5 of Audacity® software and then exported in a .mp3 and .wav file reproducible with VLC software (VideoLan, 200, VLC media player). Audio sequences were retrieved from VLC software during the test phase.

### General test procedure

Each chick was tested only once in a fully between-subjects design (in Experiments 1a and 1b, different groups of chicks were tested at different post-hatching days, see below). On the test day, chicks were transported individually from the dark incubator to the test apparatus in a closed opaque box to ensure visual naïveté.

Each chick was placed in the central area of the apparatus, confined for 1 minute within a small black plastic cage with a metal mesh front that allowed visual and acoustical access to the stimuli. This was done to promote attention to both auditory sequences before a choice could be expressed.

After the exposure phase, the cage was lifted, and chicks could freely explore the apparatus for 6 minutes (test phase). During both the exposure and the test phases, two auditory sequences were played continuously and asynchronously from the left and right speakers, alternating with 5-second silent pauses. Each 7-minute session included ∼42 sequences (21 per side/numerosity).

### Data acquisition and data analysis

We measured the time (in seconds) each chick spent in the two choice areas, and used these values to compute a preference index for the two acoustic stimuli. Time spent within a choice area was taken as an indicator of preference for the stimulus presented on that side, whereas the central zone was considered a neutral area. A choice was scored when the chick crossed the boundary of a choice area with both feet. Behaviour was recorded in real time by the experimenter using a custom Matlab function.

Raw times were converted into preference indices using the following formula:

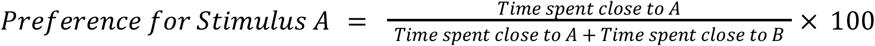

Scores above 50% indicated a preference for stimulus A, scores below 50% indicated a preference for stimulus B, and a score of 50% indicated no preference. Depending on the comparison of interest, stimulus A could represent: the sequence with more sounds (i.e. the 12-sound sequence, in Experiments 1a and b), the longer sound in Experiment 3, or the familiar sequence (Experiment 2). When relevant, we tested for an effect of test day (Experiment 1), while the effect of sex was considered only in relation to familiar preference after imprinting (Experiment 2), since sex differences in response to numerosity have been reported exclusively as a consequence of imprinting. (Lemaire et al., 2020).

All statistical analyses were conducted in R (R Core Team, 2022) within the RStudio environment (RStudio Team, 2020). Specific statistical tests are described in the results section of each experiment. In general, analyses used a significance level of α = 0.05. Effect sizes are reported as 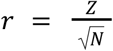 for Wilcoxon tests and η^2^ for ANOVAs. We used ANOVA to account for mixed-model designs and multiple factors; this approach is robust to violations of normality (Blanca et al., 2017). Corrections for violations of sphericity were applied when necessary, and post-hoc analyses were conducted using non-parametric tests to account for non-normal data distributions.

## EXPERIMENT 1: SPONTANEOUS PREFERENCE FOR ACOUSTIC NUMEROSITY

### Experiment 1a - No control for extensive variables (presentation rate matched)

The aim of the first experiment was to investigate whether chicks would show a spontaneous preference for a sequence containing a larger number of sounds compared to one containing fewer sounds. In this experiment, sound sequences consisted of either 4 or 12 identical sounds. Consequently, continuous variables were not controlled, as both the total duration of the sequence (including pauses) and the overall amount of sound increased with numerosity. However, because the sound structure was identical across conditions, the presentation rate (i.e., the number of sounds per minute) remained constant within each sequence.

#### Animals

A total of 136 domestic chicks (*Gallus gallus domesticus*) were tested. Of these, 68 (30 females) were examined on the day of hatching (21^st^ post fertilisation, hereafter referred to as P0), and 68 (35 females) were examined on the following day (hereafter referred to as P1).

#### Stimuli

In this experiment, a single 1000 Hz tone lasting 200 ms was used as the basic unit of the sequence. We concatenated 4 (or 12) tones with a 200 ms pause to generate a sequence of stimuli. The resulting sequences had a presentation rate of 150 bmp, mirroring the hens’ natural calls (De Tommaso et al., 2019; Kent, 1993). As we kept the inter-stimulus frequency and length of the single tones constant, the 4-sound sequences lasted 1400 ms, and the 12-sound sequence lasted 4600 ms, with a total of 800 ms and 2400 ms of played sound, respectively (Figure 3.3). During the test, the two different auditory sequences were played alternately from the left and right speakers, with a 5-s pause between successive presentations. The starting sequence and side of the apparatus assigned to each sequence were fully counterbalanced across subjects.

#### Results

We found no significant difference in preference scores between chicks tested at P0 and chicks tested at P1 (*two-tailed two-sample Wilcoxon Rank-Sum Test:* W = 2503.5, p = .343, Z = 0.256). Data were therefore collapsed across test days. Overall, chicks showed a significant preference for the larger numerosity (M = 58.46%, SEM = 4.15%; *two-tailed one-sample Wilcoxon signed-rank test*: W = 5525, p = .0346, Z = 0.181; Figure 3).

**Figure 1.**
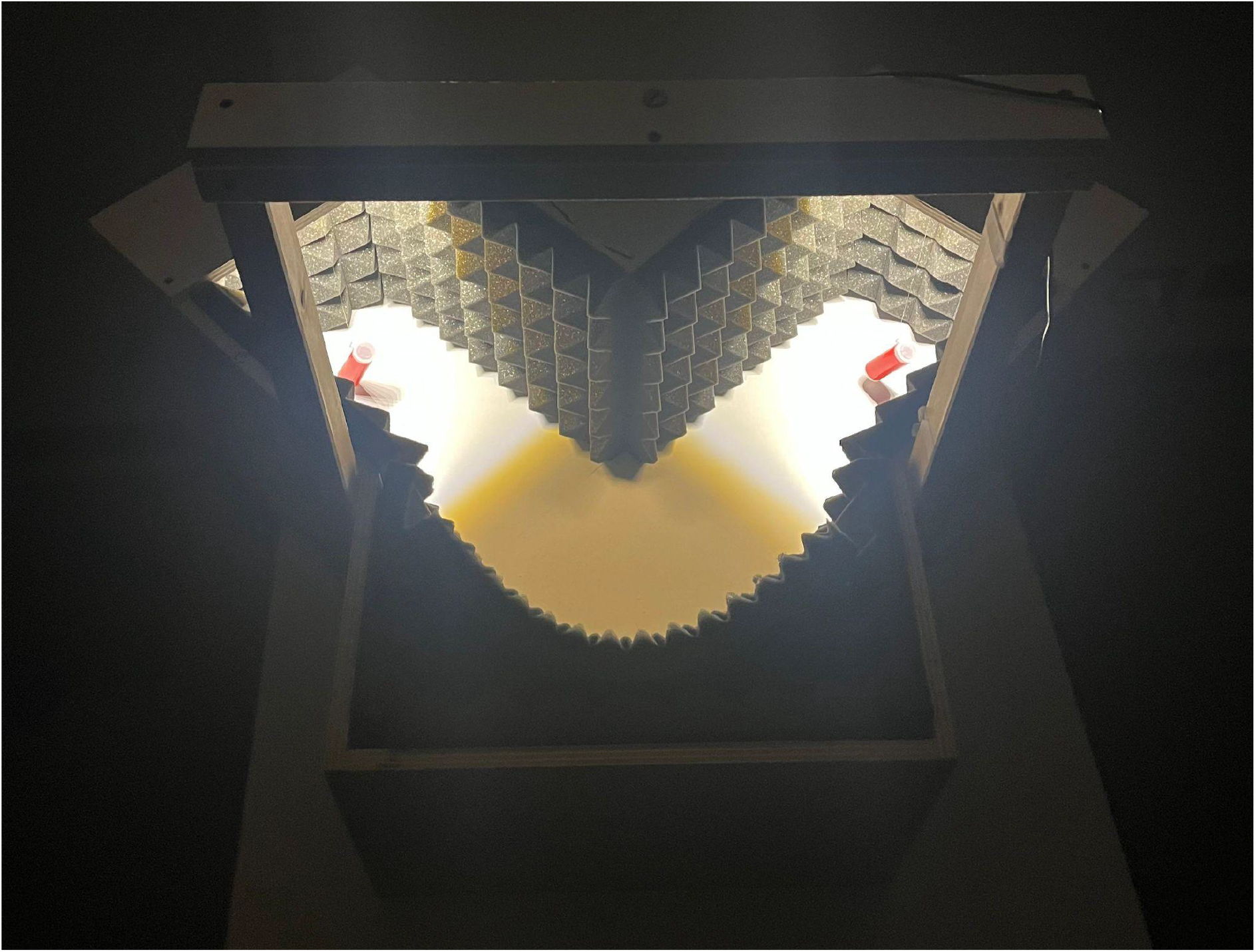
Picture of the apparatus used for all the experiments. In the picture, the apparatus used to perform all the experiments is shown. Sound-absorbing material covered the apparatus walls. Two task-irrelevant stimuli (two red cylinders) hung in front of the speakers to increase the chicks’ propensity to enter the choice areas.

**Figure 2.**
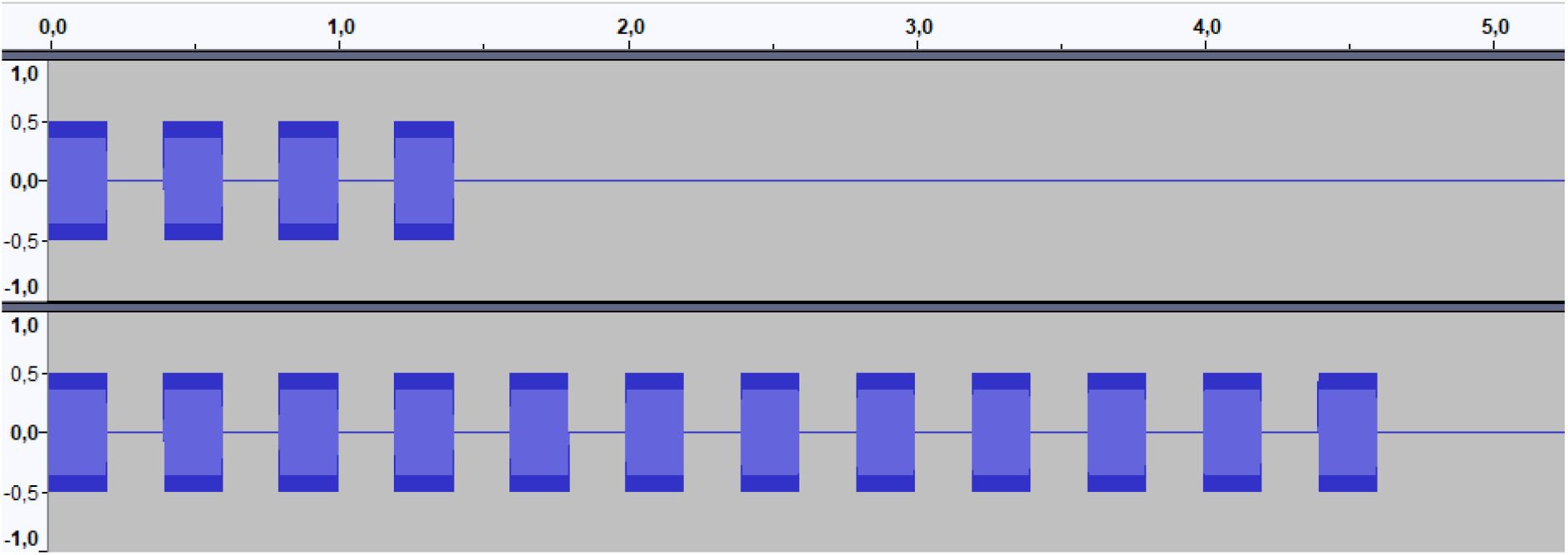
Experiment 1a stimuli. Structure of the auditory stimuli displayed in the Audacity software window, amplitude (y axis) per time (x axis, in seconds). The upper panel shows the 4-sound sequence, and the lower panel shows the 12-sound sequence

**Figure 3.**
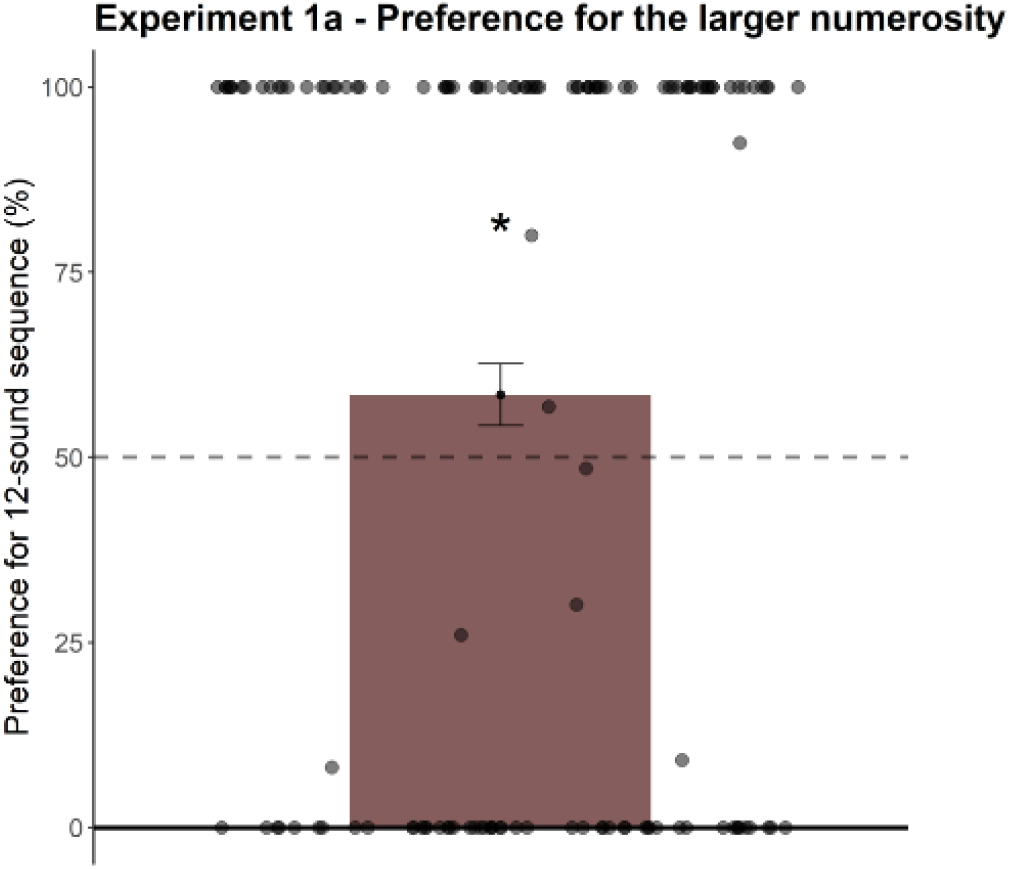
Overall preference for the 12-sound sequence across the sample. The black dotted line indicates the chance level. Data are presented as mean ± standard error of the mean; grey dots represent single data points. Statistical significance was assessed with a two-tailed one-sample Wilcoxon signed-rank test (* = p < .05)

### Experiment 1b - Control for extensive variables (presentation rate not matched)

In this experiment, the duration of the individual tones in the 4-sound sequence was increased, to equate the total sequence duration and the overall amount of sound across the two stimuli. This manipulation served to dissociate the effect of numerosity from these extensive dimensions and to evaluate their potential contribution to the preference for the 12-sound sequence observed in Experiment 1a. It should be noted, however, that this procedure necessarily introduced a difference in the presentation rate (i.e. rate of sound-to-silence alternation) between the two sequences.

#### Animals

As for the previous experiment, a total of 136 domestic chicks were tested. Of these, 68 were examined on P0 (29 females), and 68 were examined on P1 (38 females).

#### Stimuli

The 12-sound sequence was the same as Experiment 1a, while tones in the 4-sound sequences were lengthened to 600 ms with a ca 730 ms pause; therefore, the resulting sequences had a presentation rate of approximately 45 bpm. Crucially, the two sequences lasted both 4600 ms, with a total of 2400 ms of played sounds.

#### Results

As in the previous experiment, no difference was found in terms of preference score between chicks tested at P0 and at P1 (*two-tailed two-sample Wilcoxon Rank-Sum Test:* W = 2031, p = .167, Z = 0.119). Data were therefore collapsed across test days. Overall, chicks showed no significant preference for either stimulus(M = 48.79%, SEM = 4.23%; *two-tailed one-sample Wilcoxon signed-rank test*: W = 4516, p = .729, Z = 0.03).

### EXPERIMENT 2: FILIAL IMPRINTING FOR NUMEROSITY IN RESPONSE TO SOUND SEQUENCES

This experiment aimed to determine whether chicks imprinted on a sequence containing a given number of sounds would recognise and prefer that sequence over one containing a different number of sounds. To this end, from the last days of incubation onward, chicks were exposed to either a sequence of 12 sounds or a sequence of 4 sounds, and were then tested for their preference between the two. To minimise potential confounds arising from pre-existing spontaneous preferences, we used the stimuli from Experiment 1b, in which no spontaneous preference between the two sequences had been observed.

#### Animals

As for the previous experiment, a total of 136 domestic chicks were tested on P1, the day after hatching Of these, 68 (35 females) were assigned to the 4-sound imprinting condition, and 68 (31 females) to the 12-sound imprinting condition. Since the previous experiments revealed no difference in behaviour between P0 and P1, in the present experiment chicks were tested only on P1, a condition that also allowed the imprinting phase to take place after hatching.

#### Stimuli

We used the same stimuli as in Experiment 1b (Figure 4), which had not elicited any spontaneous preference in chicks. In these stimuli, sequence duration and amount of sound were balanced, allowing us to investigate the recognition of the familiar sequence independently of these two features.

**Figure 4.**
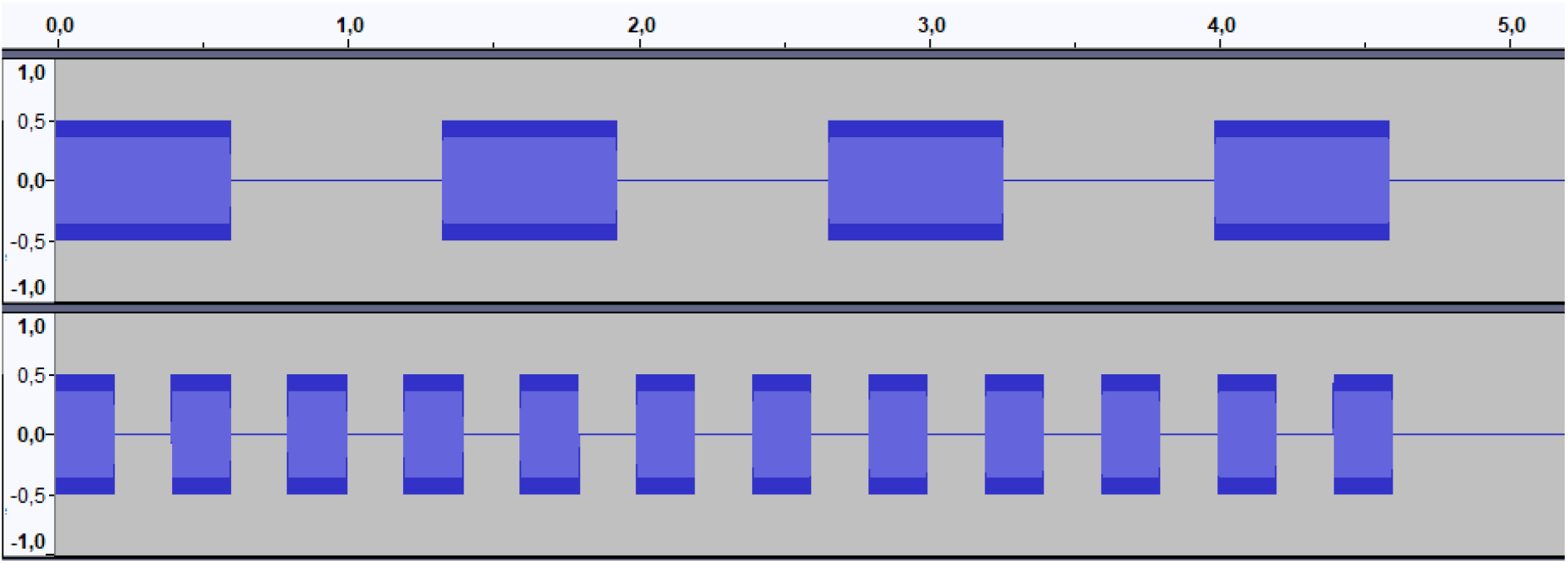
Experiment 1b stimuli. Structure of the auditory stimuli displayed in the Audacity software window, amplitude (y axis) per time (x axis). The upper panel shows the 4-sound sequence, and the lower panel shows the 12-sound sequence

**Figure 5.**
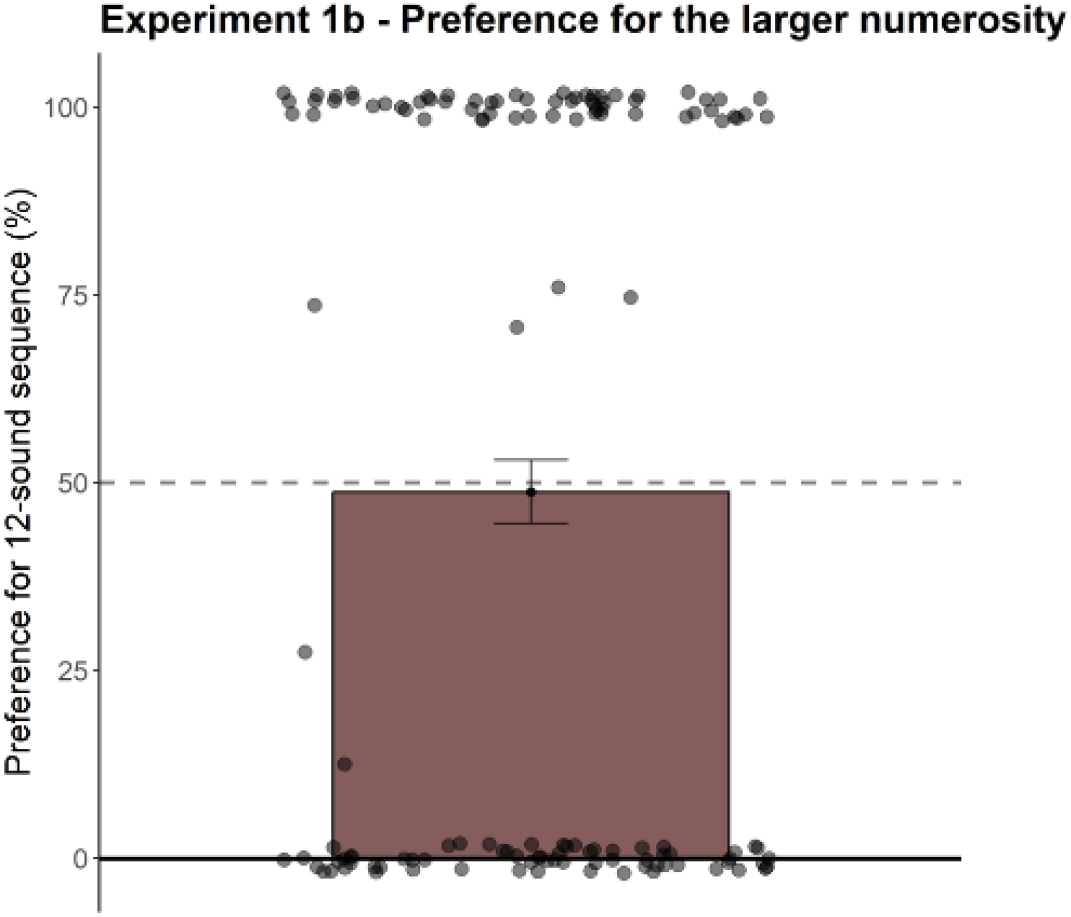
Overall preference for the 12-sound sequence across the sample. The black dotted line indicates the chance level. Data are presented as mean ± standard error of the mean; grey dots represent single data points. Statistical significance was assessed with a two-tailed one-sample Wilcoxon signed-rank test (no significant difference was found)

#### Imprinting procedure

Each batch of eggs was exposed to one of the two numerosities, in a between-subjects design. The imprinting phase started in ovo, from the 20^th^ day of development up to and including the hatching day. Eggs were incubated in the dark, as described before, and two speakers (Z130 Stereo Speakers) inside the incubator played the auditory imprinting sequence. The sound was played starting from 8 am for 8 hours a day, alternating 2 hours of stimulation with 2 hours of silence. During the 2-h stimulation period, a sound sequence composed of either 4 or 12 sounds was presented. Each sequence was repeated every 5 s for a duration of 6 min, mimicking the structure of the test procedure. Each 6-min stimulation block was followed by a 5-min silent interval. This alternation of stimulation and silence continued for the entire 2-h period. During the remaining hours, no sound was played, aiming to both reproduce an ecological situation (i.e. silence at night, since domestic chickens are diurnal animals) and to avoid the sounds becoming a background noise that the animals may ignore. Animals were tested on P1. On the morning of test day, the imprinting stimulus was played for 30 minutes as a short recall session. The test session was the same as described above.

#### Results

The two-way Imprinting Condition*Sex (2×2) ANOVA revealed a significant main effect of Imprinting Condition on preference for the familiar stimulus (F_(1,132)_ = 4.946, p = .028, η_p_^2^ = 0.036; Figure 6). Chicks imprinted on the 4-sound sequence showed a lower level of preference for the familiar sequence, compared to those imprinted on the 12-sound sequence. None of the two groups, however, showed a significant preference in either direction (chicks imprinted on the 4-sound sequence: M = 40.07%, SEM = 5.90%; *two-tailed one-sample Wilcoxon signed-rank test*: W = 951, p = .129, Z = 0.004; chicks imprinted on the 12-sound sequence M = 58.89%, SEM = 5.70%; W = 1420, p = .100; Z = 0.004). No significant effects of Sex (F_(1,132)_ = 1.521, p = .220, ηp^2^ = 0.011) or of the interaction between Imprinting Condition and Sex (F_(1,132)_ = 0.125, p = .724, ηp^2^ = 0.001) were found. Thus, chicks did not seem to have a preference for the familiar imprinting stimulus.

**Figure 6.**
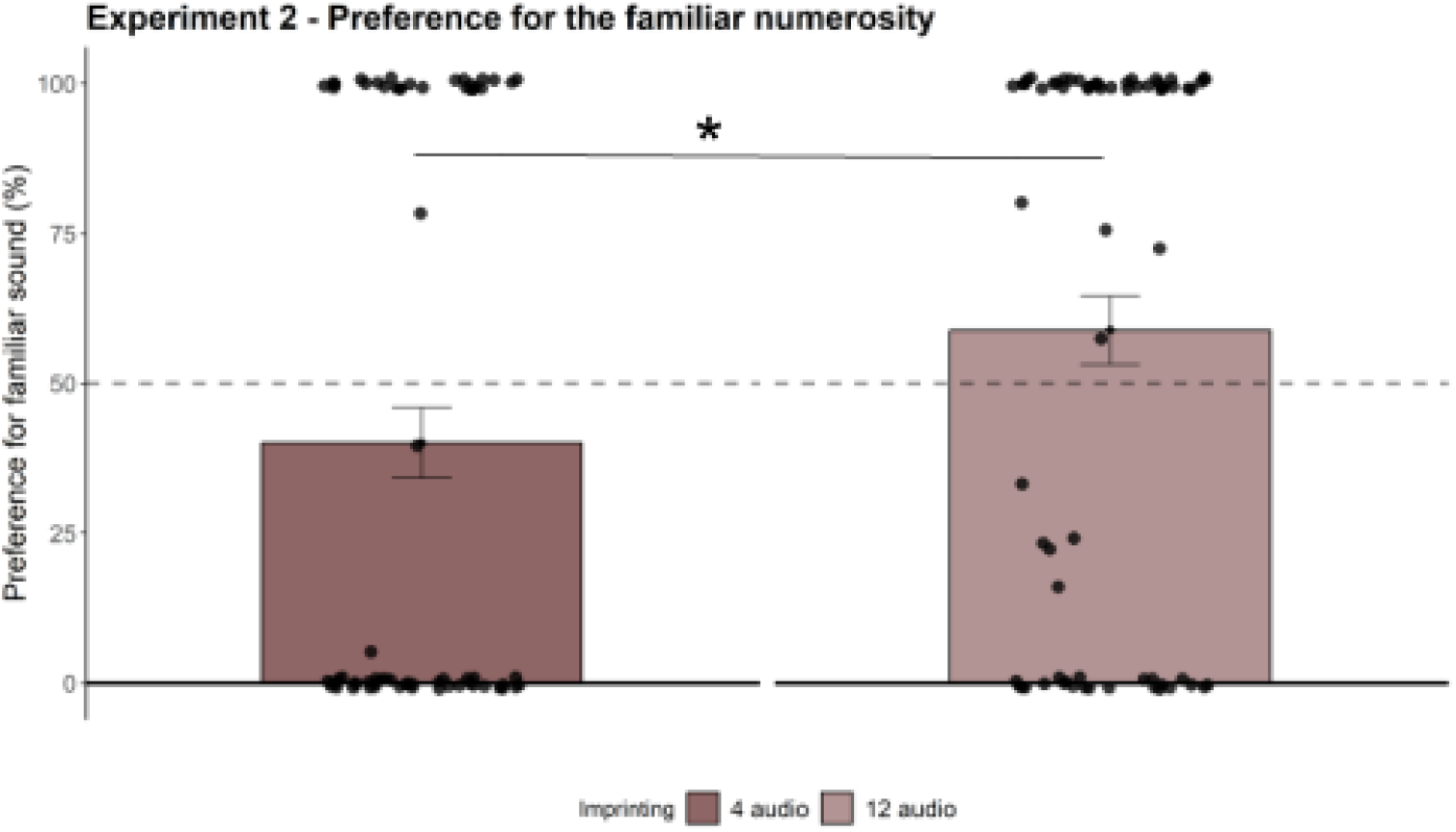
Familiar preference score in Experiment 2. Overall preference for the familiar sequence across the sample. The black dotted line indicates the chance level. Data are presented as mean ± standard error of the mean; grey dots represent single data points. Statistical significance was assessed with two-way ANOVA (* = pvalue <.05)

Nevertheless, chicks imprinted on the 4-sound sequence tended to approach the unfamiliar (12-sound) stimulus, whereas those imprinted on the 12-sound sequence showed the opposite trend, tending to prefer the familiar stimulus. In other words, both groups displayed a tendency to approach the 12-sound sequence.

To assess whether this trend was comparable across the two groups, we recalculated the preference index in terms of preference for the larger numerosity rather than for the familiar stimulus, and then compared this value between groups through a two-way (2 × 2) ANOVA. The analysis revealed no main effects of Imprinting Condition (F_(1,132)_ = 0.020, p = .888, ηp^2^ = 0.0002) or Sex (F_(1,132)_ = 0.125, p = .724, ηp^2^ = 0.001), and no significant interaction (F_(1,54)_ = 1.521, p = .220, ηp^2^ = 0.011). Data were therefore collapsed across both factors to achieve greater statistical power. Overall, chicks showed a significant preference for the larger numerosity (M = 59.41%, SEM = 4.09%; *two-tailed one-sample Wilcoxon signed-rank test*: W = 5597, p = .024, Z = 0.193; Figure 7). Therefore, imprinting appeared to induce a preference for the longer sequence, which had not emerged in Experiment 1b.

**Figure 7.**
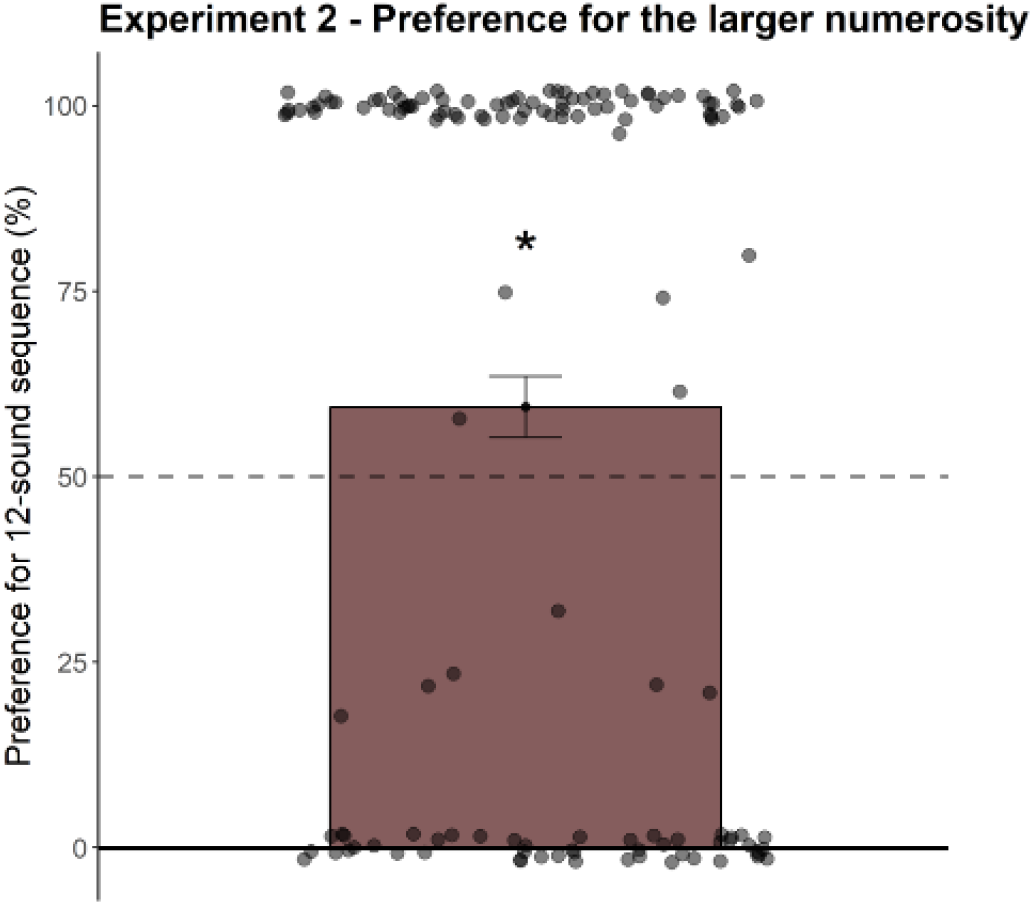
Preference Score for larger sound in Experiment 2. Overall preference for the 12-sound sequence, across the entire sample (N = 136), irrespective of imprinting condition. The black dotted line indicates the chance level. Data are presented as mean ± standard error of the mean; grey dots represent single data points. Statistical significance was assessed with a two-tailed one-sample Wilcoxon signed-rank test (* = pvalue <.05)

### EXPERIMENT 3: SPONTANEOUS PREFERENCE FOR SOUND DURATION

We conducted a third experiment to rule out the possibility that the preference for the 12-sound sequence observed in Experiment 1a was driven by a bias toward the stimulus with the greater ammount of sound, rather than by numerosity per se. In this experiment, chicks were tested only at P1 (the second day after hatching). This choice was motivated by the results of Experiment 2, which was conducted at P1 and revealed a preference for the 12-sound sequence. Testing a single age allowed us to limit the number of animals required for this control experiment.

#### Animals

A total of 68 (40 females) domestic chicks were tested on P1.

#### Stimuli

We presented two single sounds, one shorter and one longer. The sound durations have been made equal to the 4-sound sequence and the 12-sound sequence of Experiment 1a, meaning that the sounds lasted 1400 ms and 4600 ms, respectively (Figure 8). The auditory stimulus had the same pitch as in the previous experiments, ensuring consistency across conditions.

**Figure 8.**
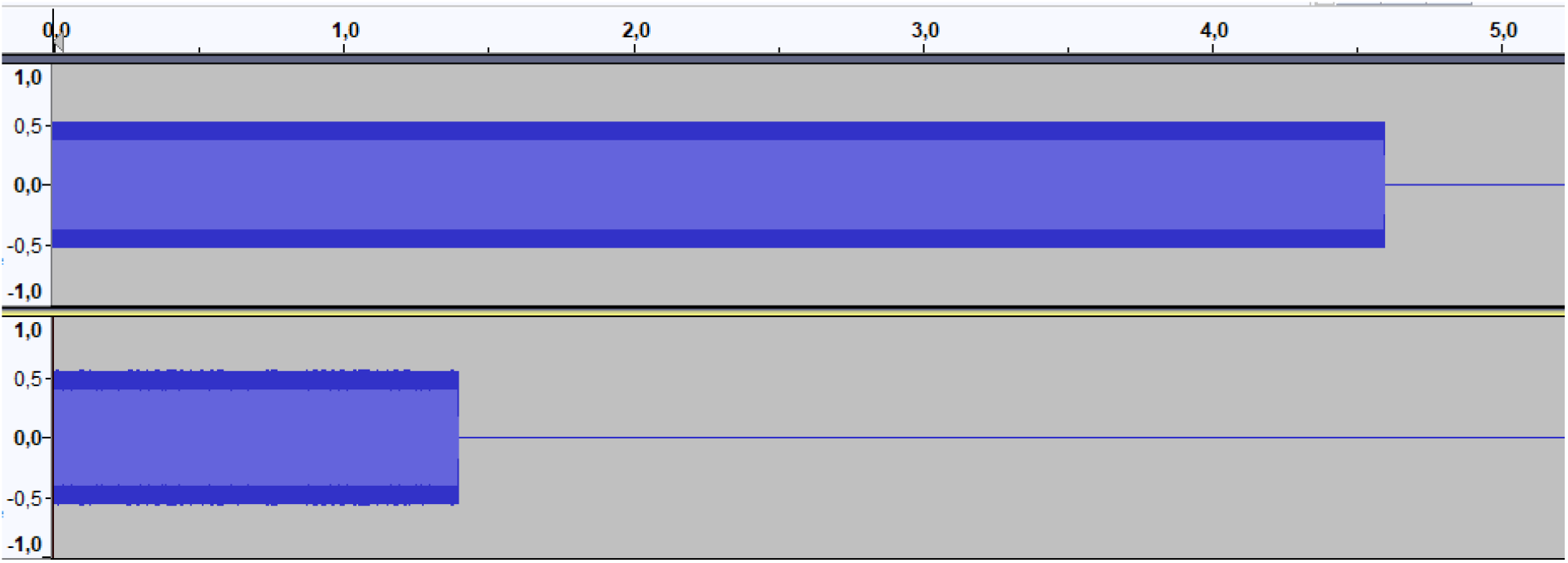
Experiment 3 stimuli. Structure of the auditory stimuli displayed in the Audacity software window, amplitude (y axis) per time (x axis). The upper panel shows the longer sound, and the lower panel shows the shorter sound

#### Results

When tested against chance level, no significant preference for the longer sequence was found (M = 39.71%, SEM = 5.98%; *two-tailed one-sample Wilcoxon signed-rank test*: W = 931.5, p = .090, Z = 0.206; Figure 9). In contrast, an opposite trend emerged: if anything, the average preference tended to be in the direction of the shorter sound.

**Figure 9.**
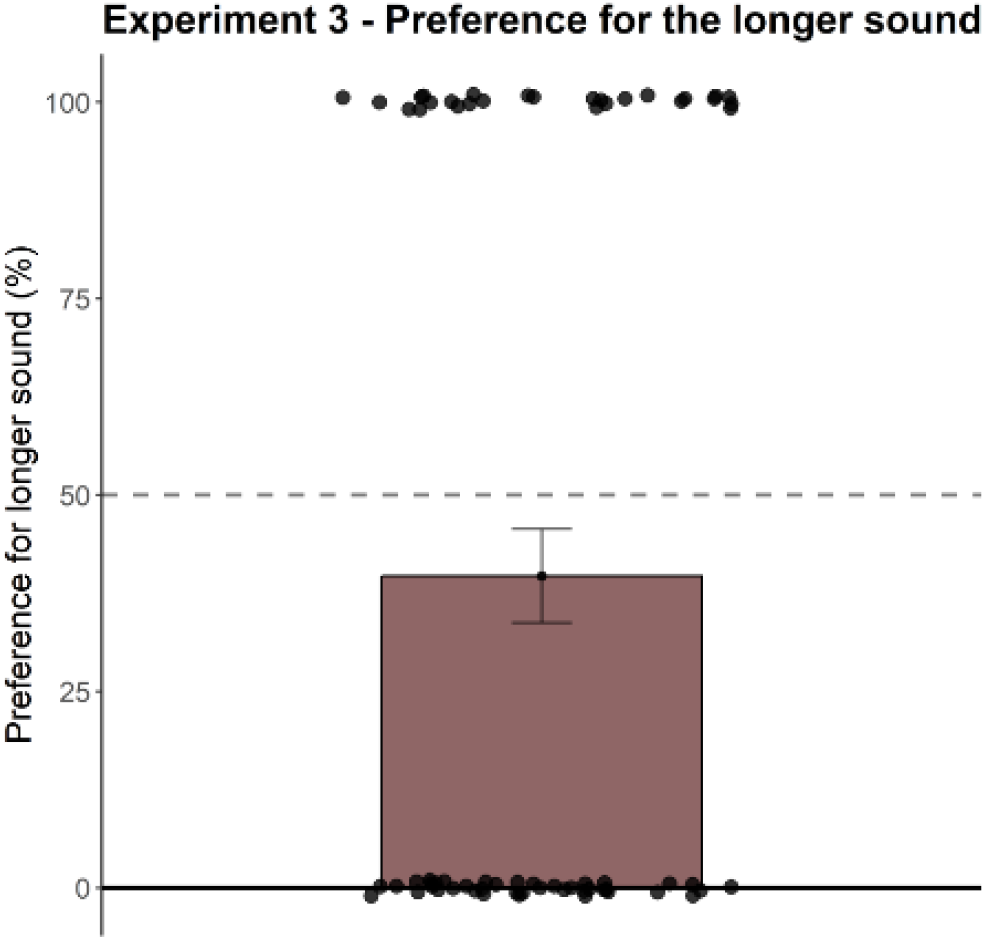
Overall preference for the 12-sound sequence across the sample. The black dotted line indicates the chance level. Data are presented as mean ± standard error of the mean; grey dots represent single data points. Statistical significance was assessed with a two-tailed one-sample Wilcoxon signed-rank test (no significant preference was found).

## DISCUSSION

In this study, we explored whether newly hatched chicks can discriminate auditory sequences that differ in the number of sounds they contain. In a first experiment (Experiment 1a), chicks were presented with sequences differing in numerosity, without manipulations of continuous acoustic dimensions. As a result, the more numerous sequence was also longer in duration and contained a greater overall amount of sound. In this context, chicks showed a significant preference for a sequence containing 12 sounds over a sequence of 4 sounds. This behaviour parallels well-established findings in the visual domain, where a preference for the set containing the larger number of imprinting objects has been repeatedly reported naïve chicks consistently approach the larger of two stimulus sets; notably, chicks reared with sets differing in numerosity later prefer the larger set at test, irrespective of the specific numerosities experienced during rearing (Rugani et al., 2010). Nonetheless, because continuous acoustic variables were not dissociated from numerosity in Experiment 1a, the observed preference cannot be unequivocally attributed to number alone, as it may also reflect sensitivity to overall duration or total sound amount. To address this possibility, in Experiment 1b we equated continuous variables across the two sequences. When duration and total sound amount were controlled, chicks no longer showed a preference for the 12-sound sequence, suggesting that additional magnitude cues were responsible for driving the preference observed in Experiment 1a. Notably, presentation rate also does not appear to have been a determining factor: despite the 12-sound sequence presentation rate in Experiment 1b being higher and more similar to natural hen calls, no preference emerged for this stimulus.

To further investigate the hypothesis that sound sequences may convey information about the numerosity of a social group, we also conducted an imprinting experiment. In Experiment 2, we tested whether chicks imprinted on a specific acoustic numerosity would later prefer to approach that numerosity at test. Consistent with previous studies on visual numerical imprinting (Lemaire et al., 2020; Rugani, Regolin, et al., 2010), chicks preferentially approached the larger numerosity rather than the one experienced during imprinting. This result is particularly noteworthy given that the same stimuli, when presented to non-imprinted chicks in Experiment 1b, had elicited no spontaneous preference for the 12-sound sequence. . Thus, exposure to the imprinting stimuli, regardless of whether these were sequences of 4 or 12 sounds, appears to have been the key factor underlying the emergence of the preference for the 12 sounds stimulus. Finally, in Experiment 3, we wanted to rule out the possibility that chicks preference for the longer sequence, in Exp. 1a, was based on the stimuli’s total length. We presented chicks with two single sounds of different duration (matching that of the sequences used in Exp. 1a), which therefore contained no numerical information. We found that chicks did not prefer one stimulus over the other.

Overall, the combined results from all the experiments suggest that the integration of both numerical and magnitude information is crucial for auditory numerical comparisons in naïve domestic chicks. Indeed, while Experiment 1b suggested that when the overall amount of sound is equalised between the two sequences, chicks do not show a preference, Experiment 3 shows that when only magnitude information (i.e. total length and overall amount of sound) is available, chicks still do not show a preference for one sound over the other. Thus, it seems that chicks’ preference for the larger sound sequence in Experiment 1a was made possible by the presence of both magnitude information (absent in Experiment 1b) *and* numerical information (absent in Experiment 3). Moreover, Experiment 2 seems to suggest that prior exposure to the test sound can lead chicks to show a preference for the more numerous set even in the absence of parallel magnitude information. Indeed, despite using the same stimuli of Experiment 1b, we found that after undergoing an imprinting phase, chicks preferred to approach the 12-sound sequence at test, regardless of the numerosity they were imprinted to.

Overall, our results support the view that both numerical and continuous magnitude information jointly contribute to discrimination in the auditory domain. The interpretation that both numerical and continuous magnitude information contribute to auditory discrimination fits with prior literature showing that the structure of the stimuli determines which cues observers use in numerical tasks. When stimuli are homogeneous, infants and non-human animals often rely on continuous variables rather than numerical information (Feigenson et al., 2002; Rugani, Regolin, et al., 2010). Conversely, heterogeneous sets promote reliance on number (Feigenson, 2005; Rugani, Regolin, et al., 2010). Because our stimuli consisted of repeated identical sounds, it is plausible that chicks in Experiment 1a exploited redundant magnitude cues, whereas no preference emerged in Experiment 1b when these cues were removed (it remains unclear, however, why, within this framework, exposure to sequences of homogeneous sounds in Experiment 2 led to the re-emergence of a preference for the longer sequence, despite the control for the overall amount of sound). A somewhat similar pattern is documented for discrimination of large numerosities (i.e., a set of elements larger than 4): human infants succeed only when supported by continuous cues (Xu & Spelke, 2000), and chicks discriminate larger ratios (e.g., 6 vs. 9) only when quantitative variables are available, whereas smaller ratios (e.g., 10 vs. 20) can be discriminated based solely on number (Rugani et al., 2013). Even in arithmetic tasks, chicks successfully sum large numerosities only when quantitative variables are not equalised (Rugani, Regolin, et al., 2011). These findings highlight that for larger sets, redundancy between numerical and continuous magnitude cues is crucial (see Rugani, 2018, for review).

The fact that we observed a preference for the larger set also fits well in the literature. Indeed, this type of preference has been extensively documented in the visual domain and is frequently used to investigate numerical abilities in domestic chicks. For example, chicks’ arithmetic skills have been demonstrated by exploiting their consistent tendency to inspect the screen concealing the larger set of imprinting objects (Rugani, Regolin, et al., 2011), and multiple imprinting studies have shown a robust preference for approaching larger groups of artificial social companions (Lemaire et al., 2020; Rugani, Regolin, et al., 2010). These tendencies are generally interpreted as adaptive, reflecting motivations such as seeking protection, social interaction, or thermoregulation, which are better provided by larger number of social companions (Pulliam, 1973; Roberts, 1996). What is novel in our study is that the chicks in Experiment 1a had not undergone any imprinting or habituation that could have enabled them to associate the test sounds with social companions. The fact that they nevertheless preferred the more numerous sound sequence suggests that, in the auditory domain, chicks may spontaneously interpret sound numerosity as a cue to group size, approaching the more numerous sequence as a way to seek proximity to a potentially larger social group.

The difference between Experiment 1b and Experiment 2, where identical stimuli elicited no preference in spontaneous choice but did in subjects pre-exposed to the acoustical sequences, may reflect important methodological differences. For instance, spontaneous choice paradigms measure an animal’s inherent predisposition to approach one stimulus over another, rather than its mere ability to discriminate between them. Accordingly, the absence of a preference does not necessarily indicate a failure to discriminate, but may instead reflect insufficient motivation to express that discrimination behaviourally. In this respect, Experiment 2 demonstrates that chicks can discriminate between the two sound sequences, even though no spontaneous preference had emerged in Experiment 1b. One possible explanation is that prior exposure during the imprinting procedure increased the social salience of the test stimuli. Because the pitch of the sounds was identical during imprinting and testing, chicks may have recognised the individual sound element on which they had been exposed, despite differences between the test sequences in numerosity, duration, and temporal rate. This familiar acoustic feature may have enhanced the motivational value of both stimuli relative to Experiment 1b, in which chicks were naïve, thereby enabling them to engage in the more demanding discrimination of the sequences even though some continuous variables had been controlled.

On the other hand, mere exposure to sounds during the imprinting phase may have had a more general activating effect, leading chicks to prefer either the higher presentation rate, which is closer to the temporal structure of hen calls (Kent, 1993), or simply the larger numerosity, independently of the specific imprinting stimulus. Previous studies have shown that pre-activation through early stimulation of various kinds determines the emergence chicks’ predispositions toward social stimuli. For instance, motor stimulation causes subsequent attraction to predisposed social cues (Johnson et al., 1985). Indeed, a wide range of early experiences, including artificial sounds, maternal calls, motor activity in darkness, handling, or exposure to salient abstract visual stimuli, facilitate the development of social predispositions (reviewed in Rosa Salva et al., 2015). Further control experiments are needed to dissociate these alternatives, for instance, by varying the type of early stimulation or using different sounds in the imprinting and test phases.

Finally, regarding why chicks in Experiment 2 showed a preference for the larger set of stimuli rather than for the familiar numerosity, we propose two explanations. The first aligns with the considerations discussed above: the preference for a larger group of companions may have strong ecological relevance that is difficult to override. It is possible that familiarity with one sound sequence was less influential than the inherent advantage of approaching a more numerous group, leading to a robust, biologically grounded tendency to prefer the larger set (in line with Rugani et al., 2010). The second explanation concerns potential methodological limitations in how the stimuli were presented during imprinting. One factor that may affect the success of imprinting on specific sequences, and thus the acquisition of numerical information, is the temporal structure of the stimulation. For example, the strength of imprinting to colored rotating objects or flashing lights is modulated by presentation rate (Chantrey, 1974). In later work, Chantrey (1976) showed that when chicks were imprinted with rapidly alternating colored stimuli, they generalised the same response to both, suggesting that quickly alternating elements can “blend” into a single perceptual representation. In the context of auditory imprinting, the time interval between individual sounds becomes critical for allowing chicks to encode a sequence as a unified but multi-element object (e.g., a 12-sound sequence). It is therefore possible that either the internal structure of our sequences (i.e., sounds presented too closely together) or the overall structure of the sound presentation during the imprinting phase (alternation every 5 seconds) was not optimal for conveying numerical information.

Further research will be necessary to determine which factors effectively facilitate numerical imprinting from auditory stimuli. Future studies could systematically manipulate sound duration, interstimulus intervals (both within and between sequences), and the timing of the imprinting procedure to clarify how temporal structure influences numerical learning.

In conclusion, this study provides new insights into numerical discrimination in domestic chicks in the auditory domain. Our findings show that chicks can discriminate between different numerosities, but their behaviour depends on the availability of redundant magnitude information and on prior experience with the stimuli, such as through imprinting. This suggests that, in the absence of explicit motivation or familiarity, chicks rely on both numerical and extensive cues to guide their approach preferences, while they could base their preferences on numerical information alone if previously familiarised with the stimuli.

